# Will climate change affect sugar beet establishment of the 21^st^ century? Insights from a simulation study using a crop emergence model

**DOI:** 10.1101/541276

**Authors:** Jay Ram Lamichhane, Julie Constantin, Jean-Noël Aubertot, Carolyne Dürr

## Abstract

Ongoing climate change has been reported to have far-reaching impact on crop development and yield in many regions of the globe including Europe. However, little is known about the potential impact of climate change on specific stages of the crop cycle including crop establishment, although it is a crucial stage of the annual crop cycles. For the first time, we performed a simulation study to pinpoint how sugar beet sowing conditions of the next eight decades will be altered under future climate change and if these variations will affect sowing dates, germination and emergence as well as bolting rates of this crop. We chose Northern France as an important study site, representative of sugar beet growing basin in Northern Europe. Sugar beet emergence simulations were performed for a period between 2020 and 2100, taking into account five sowing dates (mid-February, 1^st^ March, mid-March, 1^st^ April and mid-April). Soil water contents and temperatures in the 0-10 cm soil horizon were first simulated with the STICS soil-crop model using the most pessimistic IPCC scenario (RCP 8.5) to feed the SIMPLE crop emergence model. We also evaluated the probability of field access for the earlier sowings, based on the amount of cumulated rainfall during February and March. When analyzed by sowing date and for successive 20-year period from 2020 to 2100, there was a significant increase in seedbed temperatures by 2°C after 2060 while no change in cumulative rainfall was found before and after sowings, compared with the past. Emergence rate was generally higher for 2081-2100, while time to reach the maximum emergence rate decreased by about one week, compared with other periods, due to higher average seedbed temperatures. The rate of non-germinated seeds decreased, especially for the earlier sowing dates, but the frequency of non-emergence due to water stress increased after 2060 for all sowing dates, including the mid-February sowing. Bolting remains a risk for sowings before mid-March although this risk will be markedly decreased after 2060. The changes in seedbed conditions will be significant after 2060 in terms of temperatures. However, the possibility of field access will be a main limiting factor for earlier sowings, as no significant changes in cumulative rainfall, compared with the past, will occur under future climate change. When field access is not a constraint, an anticipation of the sowing date, compared to the currently practiced sowing (i.e. mid-March), will lead to decreased risks for the sugar beet crop establishment and bolting. The use of future climate scenarios coupled with a crop model allows a precise insight into the future sowing conditions, and provide helpful information to better project future farming systems.

## 1. Introduction

Seed germination and seedling emergence are critical phases of a crop cycle that affect the success or failure of any crop establishment (Villalobos et al. 2016). These early phases of crop cycle are affected by several biotic and abiotic factors that may reduce seed germination and seedling emergence rates (Lamichhane et al. 2018). More specifically to abiotic stresses, many factors including seedbed water content, temperature, and the frequency and quantity of cumulated rainfall profoundly impact crop establishment (Gallardo-Carrera et al. 2007; Constantin et al. 2015; Dürr et al. 2016). Several studies reported that climate change will result in increased mean temperature and higher precipitation variability in many regions of the globe including Europe (Pendergrass et al. 2017; Kjellström et al. 2018). The effects of ongoing climate change on crop yield have been extensively studied (Lobell et al. 2008; Challinor et al. 2014). For instance, climate change from 1980 to 2008 has resulted in reduced global production of maize by 3.8% and wheat by 5.5% compared with a counterfactual without climate change (Lobell et al. 2008). A recent meta-analysis (Challinor et al. 2014) -- based on 1,700 published simulation studies on climate change impacts on yields and adaptation -- showed that without adaptation, there will be losses in production for wheat, rice and maize in both temperate and tropical regions by 2 °C of local warming.

While many studies assessed the impact of climate change on crop yields, there is less detailed information about the potential effect of climate change on crop establishment, although it is a crucial stage for annual crops. This prevents stakeholders from mobilizing adaptation strategies that may be helpful to attenuate climate change effects. Rather small adjustments (e.g. changes in varieties, sowing date and density, tillage or tactical pest management) in contrast to more systemic changes (e.g. changes in crop sequences; moving from dryland to irrigated systems or from spring to autumn sowings), may ensure successful crop establishment with positive impacts on crop yield (reviewed by Lamichhane et al. 2018). Indeed, either a lack or an excess of soil temperature, water content or rainfall may be detrimental to crop establishment. For example, if no precipitation occurs after sowing the seed imbibition process is hindered and seeds cannot germinate. In contrast, if heavy rainfall occurs following sowing soil crusting will occur preventing seedlings from being emerged (Gallardo-Carrera et al. 2007). Spring crops are more sensitive to seedbed sowing conditions than winter crops. The risk of poor crop establishment is higher for these crops also because most of them are not able to compensate a lower plant density via tillering or ramification during their development.

Sugar beet (*Beta vulgaris* L.) is a typical example of spring crop highly sensitive to seedbed sowing conditions. In Northern Europe, these conditions are frequently unfavorable, with low temperatures, heavy rainfall followed by dry periods leading to soil surface crusting on loamy soils (Dürr and Boiffin 1995). Sugar beet growers have to optimize sowing dates and seedbed preparations to ensure successful crop establishment. In addition, sugar beet is subject to bolting, if cold temperatures occur following early sowings (Longden et al. 1975; Milford et al. 2010), with negative impact on its yield and volunteer plant’s control. Simulation studies are useful to help decision-making process. Exploration of adaptation strategies to climate change using process-based models allows crop-level evaluation and adaptation of existing cropping systems (Challinor et al. 2014). While numerous crop models have been developed and used to facilitate decision-making during the crop development phase, only few models focus on the crop establishment phase.

The objective of this simulation study was to pinpoint whether sowing conditions of the next decades (2020-2100) will be altered under climate change and if these variations will affect germination and emergence, as well as bolting rates of sugar beet in Northern Europe. A total of 405 sugar beet emergence simulations were performed taking into account five sowing dates. To this aim, we first mobilized the STICS soil-crop model (Brisson et al. 1998, 2003) to generate soil water content and temperature in the seedbed (0–10 cm) using the most pessimistic IPCC scenario. We then used the data obtained as input variables to feed the SIMPLE crop emergence model (Dürr et al. 2001; Constantin et al. 2015). The emergence courses and final rates, and causes of no-seedling emergence are analyzed and discussed. In addition, the possibility of field access was evaluated comparing historical records in relation to future climatic conditions.

## 2. Materials and methods

### 2.1. Description of the SIMPLE crop emergence model

A comprehensive description including the functioning of the SIMPLE model and the list of equations and input variables has been previously provided (Dürr et al. 2001). Briefly, the model predicts the germination and emergence process and their final rates in relation to environmental conditions during sowing. The model has previously been parameterized for a number of crop species -- including wheat, sugar beet, flax, mustard, French bean, oilseed rape (Dürr et al. 2001; Dorsainvil et al. 2005; Moreau-Valancogne et al. 2008; Dürr et al. 2016), and a plant model *Medicago truncatula* (Brunel et al. 2009) – which allows to compare a range of plant species using the same set of parameters (Gardarin et al. 2016).

SIMPLE creates 3D representations of seedbeds with sowing depth distribution and the size, number, and position of soil aggregates as input variables. Daily soil temperature and soil water potential in several layers are also used as input variables for simulations, along with plant characteristics for germination and seedling growth. The model predicts germination and emergence, seed by seed, at daily intervals. The time required for germination of the seed i is chosen at random in the distribution of thermal times that characterizes the seed lot used. Cumulative thermal time from sowing is calculated above the base temperature (Tb) for germination, provided that the soil water content at the seed sowing depth is above the base water potential (Ψb). The Tb and Ψb thresholds for germination are input variables. If seed i has not germinated after a given time (usually fixed at 30 days for the simulation), the model considers that the seed will never germinate. If the seed germinates, then a seedling grows from the seed. Time is expressed as thermal time using the Tb value. To better include the effect of early water stress on seedling growth, we added a water stress function to the SIMPLE model, which reduces emergence after germination (Constantin et al. 2015). With this function, the fate of seedlings is determined by considering soil water potential in the soil layer in which the radicle grows in the two days following germination. During this period, if soil water potential is lower than Ψb, the seedling does not emerge and dies the following day. If this is not the case, the time it takes for the seedling to reach the soil surface after germination is calculated by SIMPLE based on the seed’s sowing depth, the length of the pathway the shoot takes through the aggregates, and the shoot’s elongation function, whose parameters are input variables. The probability of the seedlings remaining blocked under aggregates depends on the size and position of the clods in the seedbed, i.e. laying on the surface or below it. Soil surface crusting depends on cumulative rainfall after sowing; a proportion of seedlings remain blocked under the crust depending on daily crust water content (no seedlings are blocked if the crust is wet). Simulations at the individual seed level are run 1000 times to predict the emergence rate and final emergence percentage. The causes of non-emergence simulated by SIMPLE are (i) non-germination, (ii) death of seedlings caused by water stress after germination and (iii) mechanical obstacles (clods or a soil crust). The SIMPLE model does not consider biotic stresses, such as pests and diseases or the effect of high temperatures, which could inhibit germination or cause young seedling death.

Bolting risk is represented by a function that was not initially presented in the seminal paper describing the SIMPLE model (Dürr et al, 2001). This function was derived from Longden et al (1975) and the probability of bolting μb is calculated as:

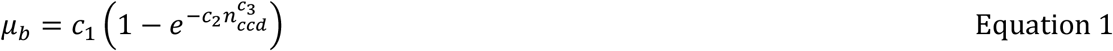

where c_1_, c_2_, c_3_ dimensionless coefficients (Table 1); n_ccd_ is the number of cumulative cold days from sowing to the end of June and is calculated as follows:

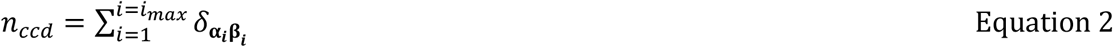

where *i* is a daily index ranging from 1 (sowing day) to *i*_*max*_ (day corresponding to the end of June) and 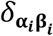 is the Kronecker symbol with:

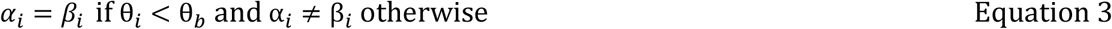

where θ_*i*_ is the maximum daily temperature at 2 m, and θ_*b*_ is the maximum threshold air temperature to define whether a given day is considered as cold or not with regard to bolting **(Table 1)**.

**Table 1.**
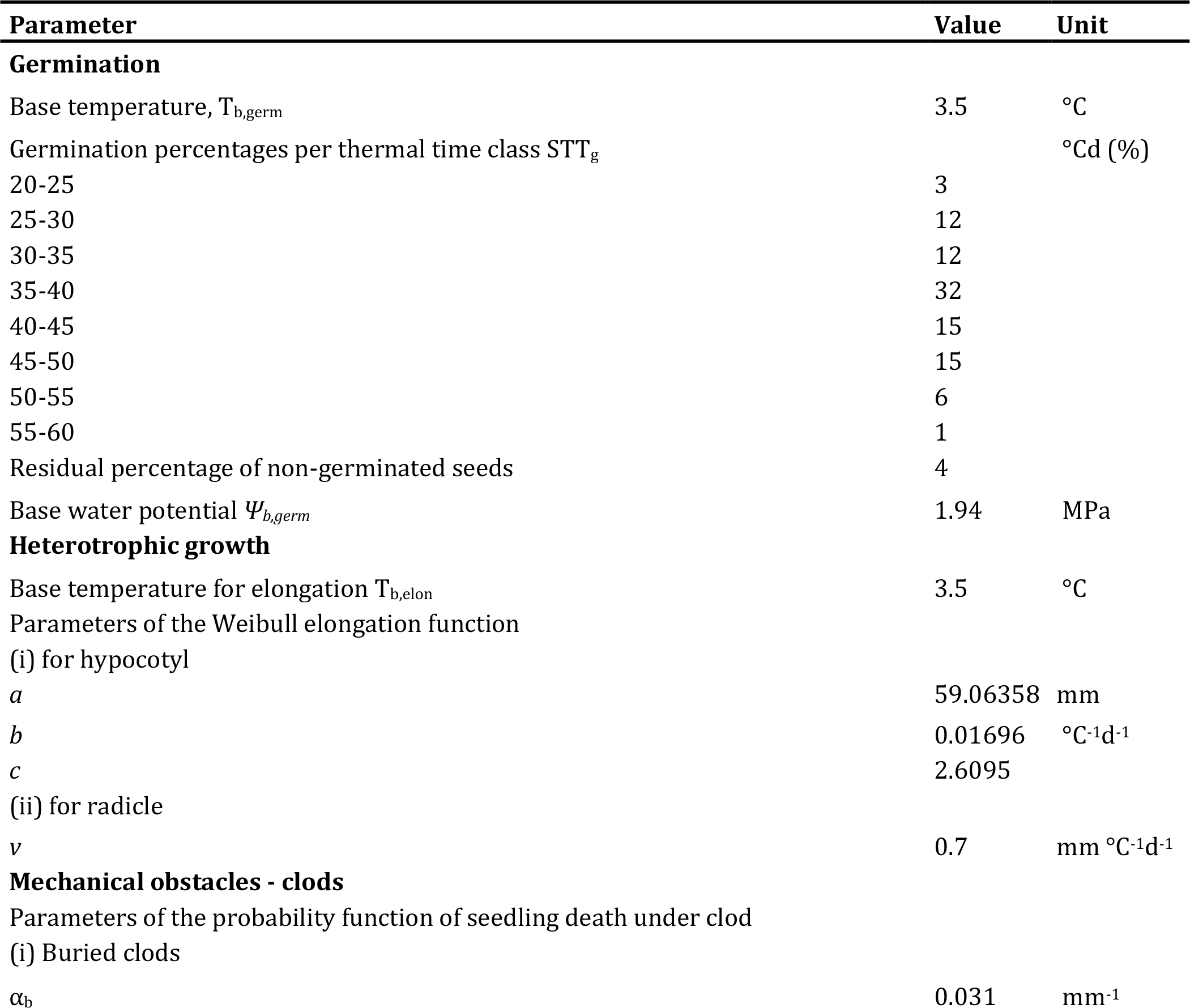
Values of the input variables of SIMPLE for sugar beet used in this study

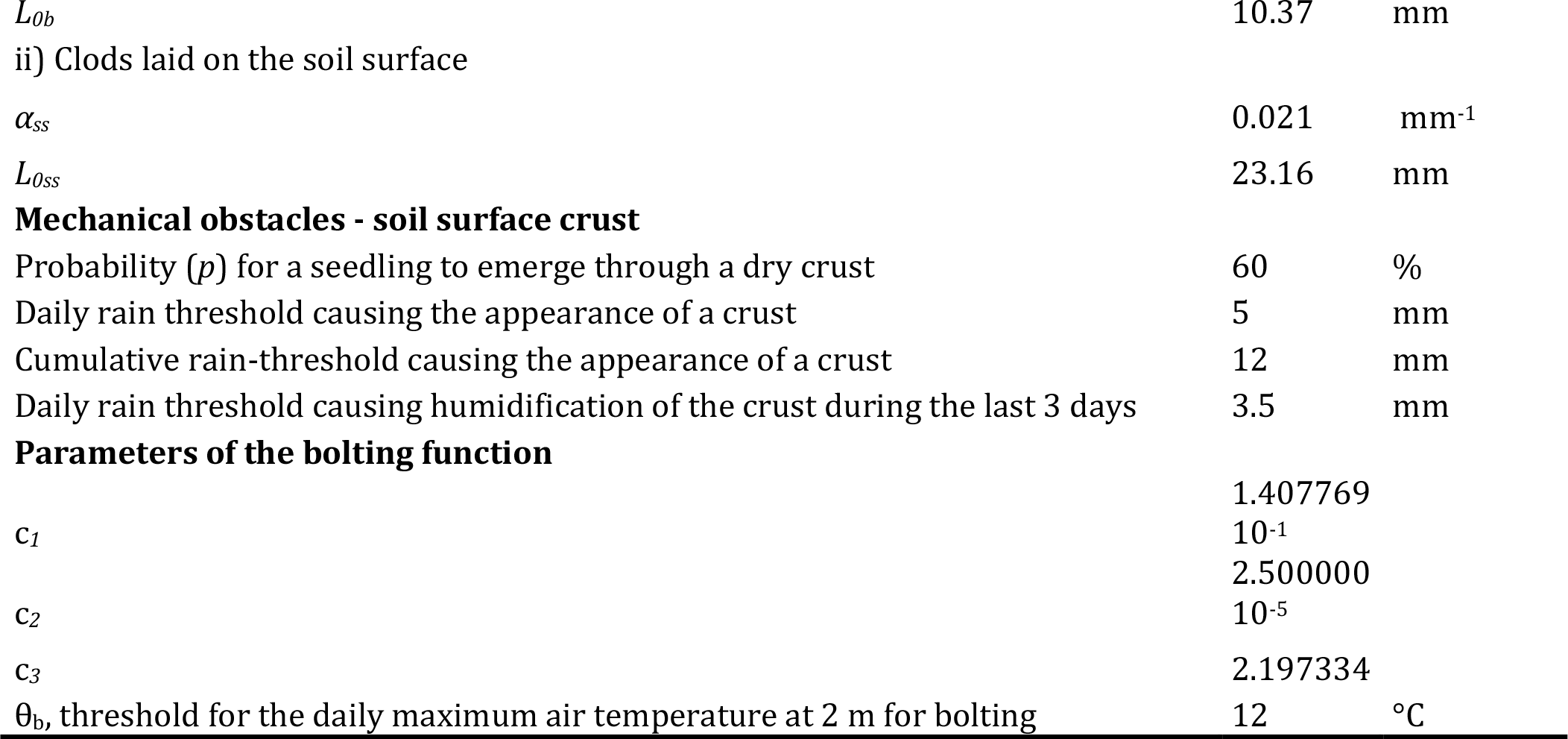

More recent studies (Fauchère et al 2003; Milford 2010) suggested that devernalization can occur if plants are exposed to high temperatures during a specific period of the crop cycle. Based on this information, we analyzed the number of days with Tmax >25°C between 60 to 120 days post sowing (dps). Finally, we established that if this number was >7, the potential risk of having bolted plants became zero.

### 2.2. Climate scenarios and simulations of the seedbed climate

We used the RCP 8.5 emission scenario to generate soil temperature and water content of the seedbed using the STICS soil-crop model (Brisson et al. 2003). This model daily simulates soil water contents and temperatures, according to daily weather and soil characteristics. Variations in soil moisture were predicted using STICS at 0-2, 2-4, 4-6 and 6-10 cm. We selected Estrées-Mons (49°52′44″N 3°00′27″E), located in the typical sugar beet growing regions of Northern France, as study site. We chose Northern France as representative sugar beet growing basin of Northern Europe. The soil type considered had the following soil granulometry and chemical characteristics at the 0-30 cm soil horizon: 0.197 g.g^−1^ clay, 0.747 g.g^−1^ silt and 0.056 g.g^−1^ sand; 0 g.g^−1^ CaCO3, 0.095 g.g^−1^ C, .001 g.g^−1^N, C/N ratio 9.3, and pH 7.7.

Four weather and soil parameters were analyzed for the year 2020-2100: average soil temperature at sowing, average soil and maximum air temperature 30 dps, and cumulated rainfall 30 dps. The average weather data of the last 19 years (2000-2018) registered at the weather station of the study area were calculated to compare the trend with the simulated weather data of the next 81 years.

### 2.3. Sugar beet sowing scenarios

Values of plant input variables of SIMPLE for sugar beet crop are reported in **Table 1**. The seedbed considered in this study is typical of that prepared by growers and was characterized by 15-25% of soil aggregates >20mm in diameter and 70-85% of its aggregates having <20mm in diameter. The simulated sowing depths were 2.5 ± 0.4 cm.

A total of 405 sugar beet emergence simulations was performed for a period between 2020 and 2100, taking into account five sowing dates: mid-February, 1^st^ March, mid-March, 1^st^ April, and mid-April. Farmers in Northern France most often practice mid-March sowing of sugar beet crop but we included both earlier (mid-February and 1^st^ March) and late (1^st^ April and mid-April) sowing dates taking into account a possible shift in future sowing dates due to climate change.

### 2.4. Analysis of simulation results

Climatic data were pooled and analyzed by sowing date and 20-year period (2000-2018 for the past and 2020-2040, 2041-2060, 2061-2080, and 2081-2100 for the future). When data were analyzed by sowing date, the 100 years were treated as replicates. When data were analyzed by 20-year period, the 20 years × five sowing dates (i.e. 100) were treated as replicates. ANOVA was used to determine the potential effect of sowing dates and periods, and their interaction on the four average weather and soil parameters mentioned above.

The variability of germination and emergence rates and their duration was analyzed by establishing three classes of rate or duration, expressed as the frequency of each class over the 20-year period for germination and emergence rates, and their duration. For germination rate, thresholds were poor germination when germination rate was <75% and sufficient germination above 75%. For emergence rate, thresholds were poor emergence when the emergence rate was <50%, and sufficient over 50%. For germination duration, thresholds were low duration when the number of days required to reach maximum germination (NGmax) was < 14 days and high when NGmax was >14 days. For emergence duration, thresholds were low duration when the number of days required to reach maximum emergence (NEmax) was < 28 days, and high when NEmax was >28 days. The frequency of poor germination (<75%) and emergence (<50%) rates as well as high NGmax (>14 days) and NEmax (>28 days) duration were analyzed as they could lead to crop emergence failure and potential re-sowing.

The variability of causes of non-emergence was analyzed by establishing two classes of seed and seedling mortality rates for each mortality cause. For non-germination, the two classes were low with <25% and high with >25% non-germinating seeds. For seedling mortality due to clod, crust and drought, the two classes were low with <15%, and high with >15% of seedling mortality. Frequency of high risks of non-germination (>25%) and seedling mortality due to clod, crust, and drought (each >15%) cases are presented for the same reason as described above.

The variability of bolting rates was presented as the average predicted percentages of bolted plants over the 20-year periods. This variability was also analyzed by establishing three classes of bolting rates : <0.5%, 0.5-1%, and >1% rate.

To determine significant effects on germination, emergence and bolting rates, and duration as well as on causes of non-emergence, in addition to the same statistical analysis performed for weather data (i.e. only by sowing date and 20-year period pooling all the data), we also analyzed the data by sowing date for each 20-year period separately (hereafter referred to as period). All statistical analyses were conducted using software R (Hothorn and Everitt 2009).

### 2.5. Technical feasibility of sowing

An earlier sowing than the currently practiced sowing (mid-March) may be possible under future climate change. This shift in sowing date however depends on field access for sowing. We determined whether farmers will have technical possibility for sowing for the simulated sowing dates and years using two following approaches.

i. based on a past historical data set (1987-2005), we observed a correlation between the quantity of total cumulative rainfall during the sowing in March and the percentage of sugar beet surface sown in France at the end of this month **(Figure 1)**. We then compared these past observations with the predicted cumulative future rainfall of the same months (i.e. February and March).
ii. we supposed that >1 mm rainfall on sowing day will not technically allow field access for farmers for the sowings in February and March because the soil surface is wet and evapotranspiration low. Based on this, we calculated the frequency of days >1 mm rainfall for February and March.

**Figure 1.**
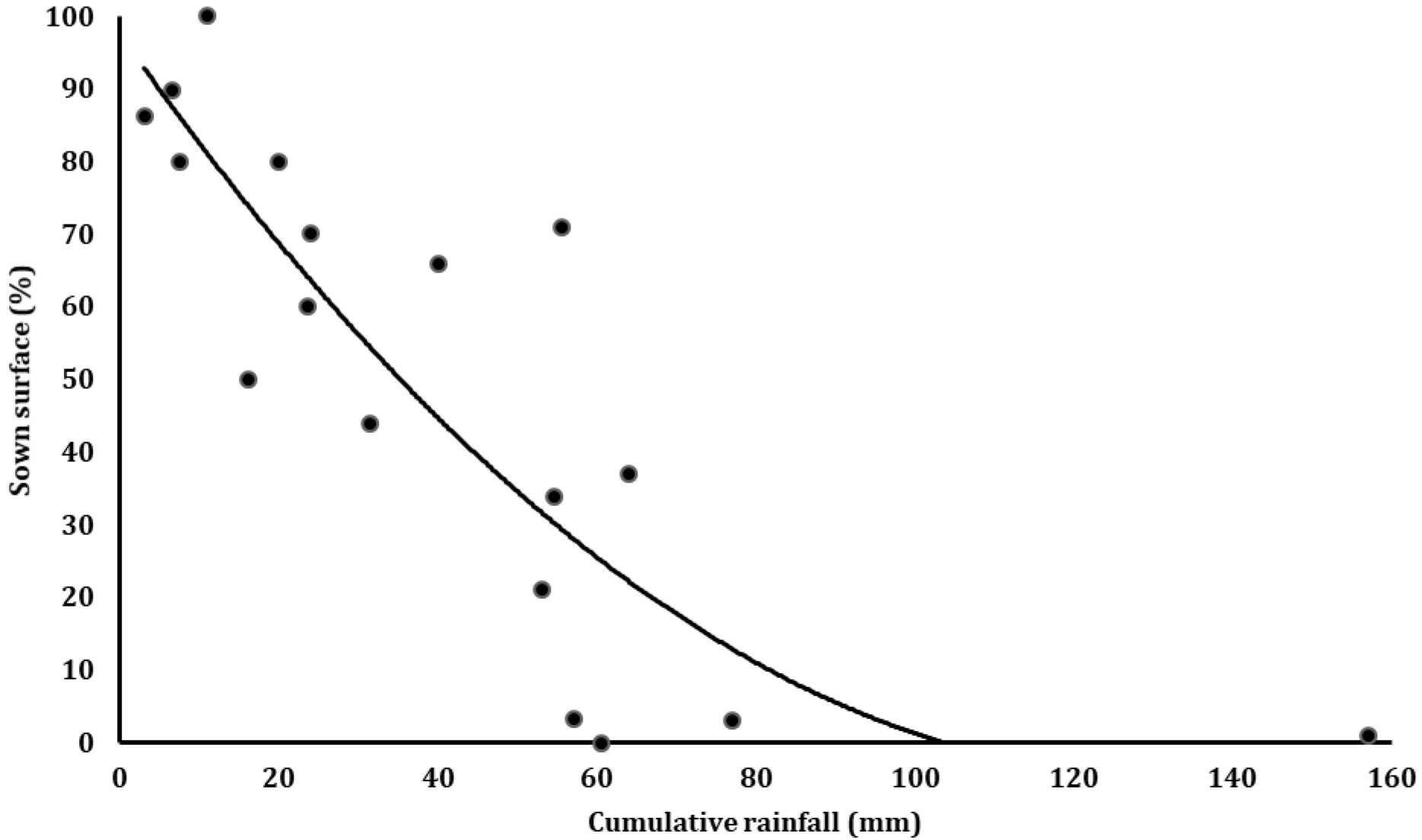
Relationship between cumulative rainfall (measured at Estrées-Mons; 49°52′44″N 3°00′27″E) and percentage of sown surface recorded at the end of March across the sugar beet growing area in France (1987-2005).

## 3. Results

### 3.1. Sowing conditions and their variability under future scenario

When analyzed by sowing date, differences between the average soil temperature at sowing, average soil and maximum air temperature 30 dps were statistically significant (P < 0.001) **(Table 2)**. When analyzed by period of time, all average weather values related to temperature increased with time with statistically significant differences (P < 0.001). In contrast, no differences statistically significant were found for average rainfall 30 dps (P = 0.220).

**Table 2.**
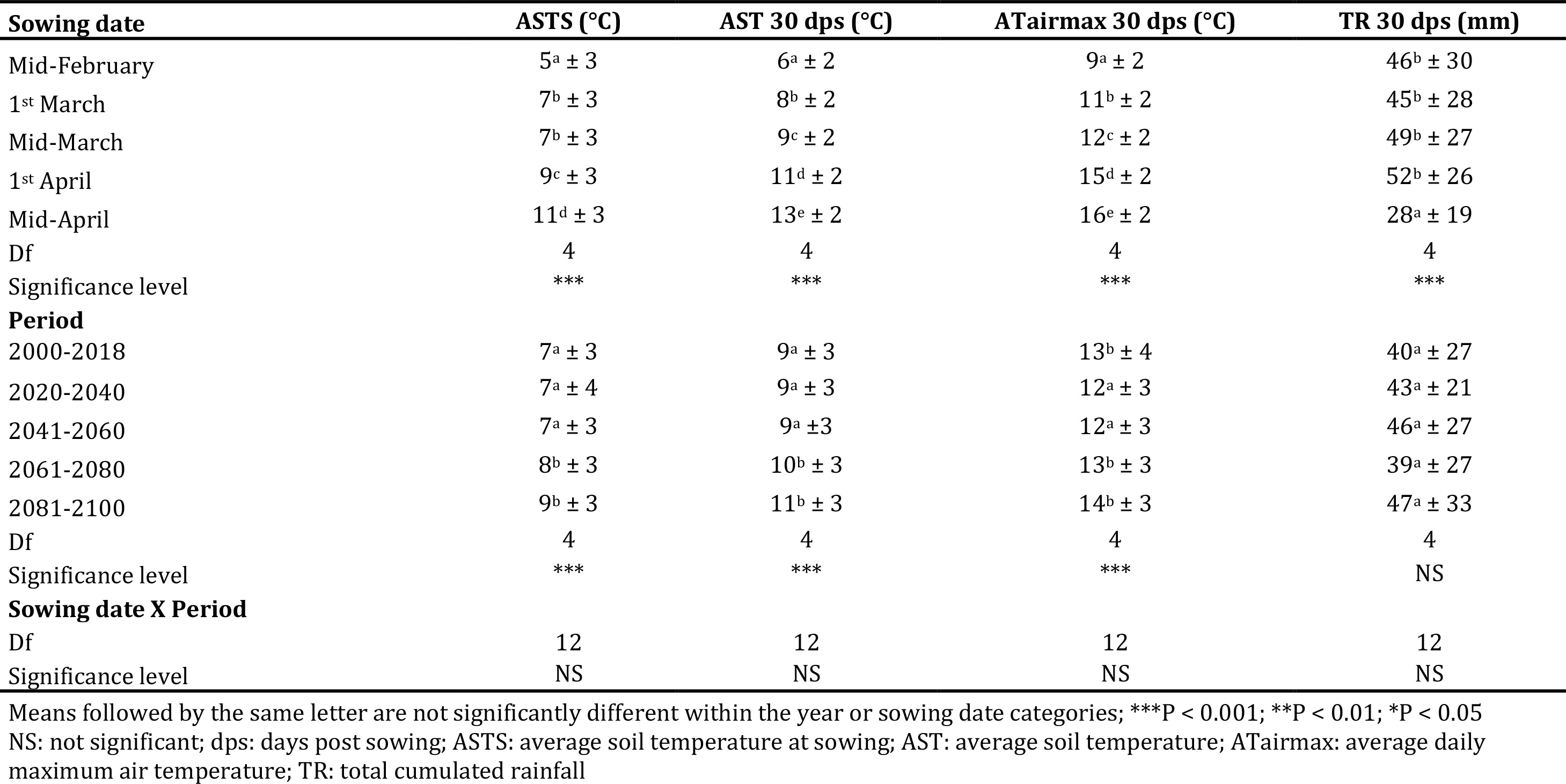
Differences among weather data (means ± standard deviation) of the study site when analyzed by sowing date, 20-year period and their interaction

As expected, average soil temperature at sowing increased with later sowing dates. The trend was similar for average soil and maximum air temperatures 30 dps. Overall, average soil temperature at sowing ranged between 5 °C for the mid-February sowing to 11 °C for the mid-April sowing, while average soil temperature 30 dps ranged from 6 °C for the mid-February sowing to 13 °C for the mid-April sowing. Also average maximum air temperature 30 dps was the lowest (9 °C) for the mid-February sowing while it was the highest (16 °C) for the mid-April sowing. In contrast to the three temperature factors, average rainfall 30 dps did not follow the same pattern: it was high (45-52 mm) for the first four sowing dates with no significant differences, and then decreased drastically for the mid-April sowing (28 mm).

When analyzed by period, average soil sowing day temperature was the lowest (7 °C) for the 2020-2040 and 2041-2060 periods, and increased progressively for the 2061-2080 and 2081-2100 periods by 8 and 9 °C, respectively. The trend was the same also for average soil temperature 30 dps, which ranged from 9 °C for the 2020-2040 period to 11 °C for the 2081-2100 period. These differences were significant between the first two and the last two periods. These changes became significant after 2060. In contrast, mean average maximum air temperature 30 dps varied over periods but with no regular increase. Mean cumulated rainfall 30 dps ranged from 39 to 47 mm, with a high variability between individual years, but without any significant differences until 2100. There was no significant effect of the sowing date × period interaction on any of the analyzed weather data **(Table 2)**.

### 3.2. Emergence rate, duration and frequency

Year-to-year emergence rate variability for all five sowing dates is described in **Figure 2**. Results on the effect of sowing date for each period and their interaction on emergence rate, duration and frequency are reported in **Table 3**. Results on the effect of sowing date and period separately, and their interaction on emergence rate and duration are presented in **Supplementary Table 1**.

**Figure. 2.**
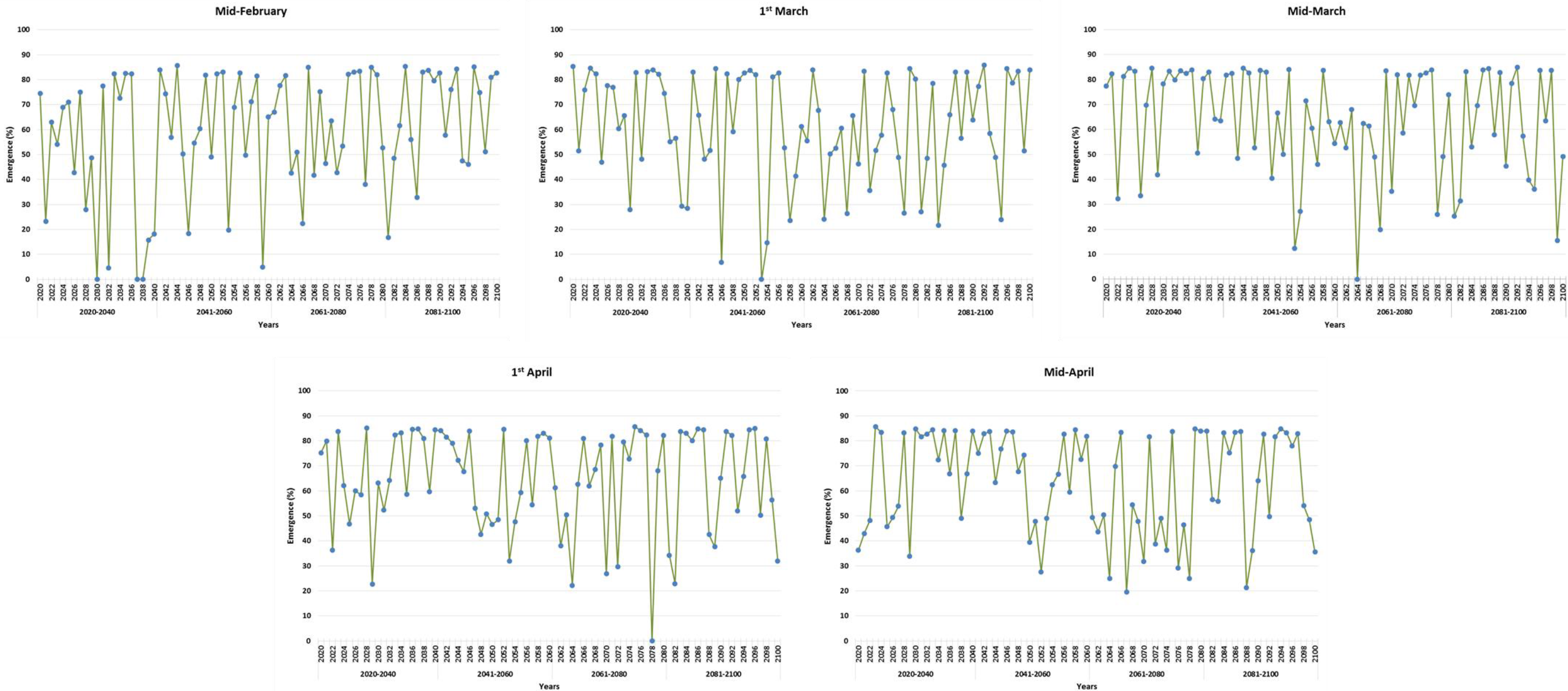
Year-to-year emergence rate variability of sugar beet by sowing date under future climate scenario

**Table 3.**
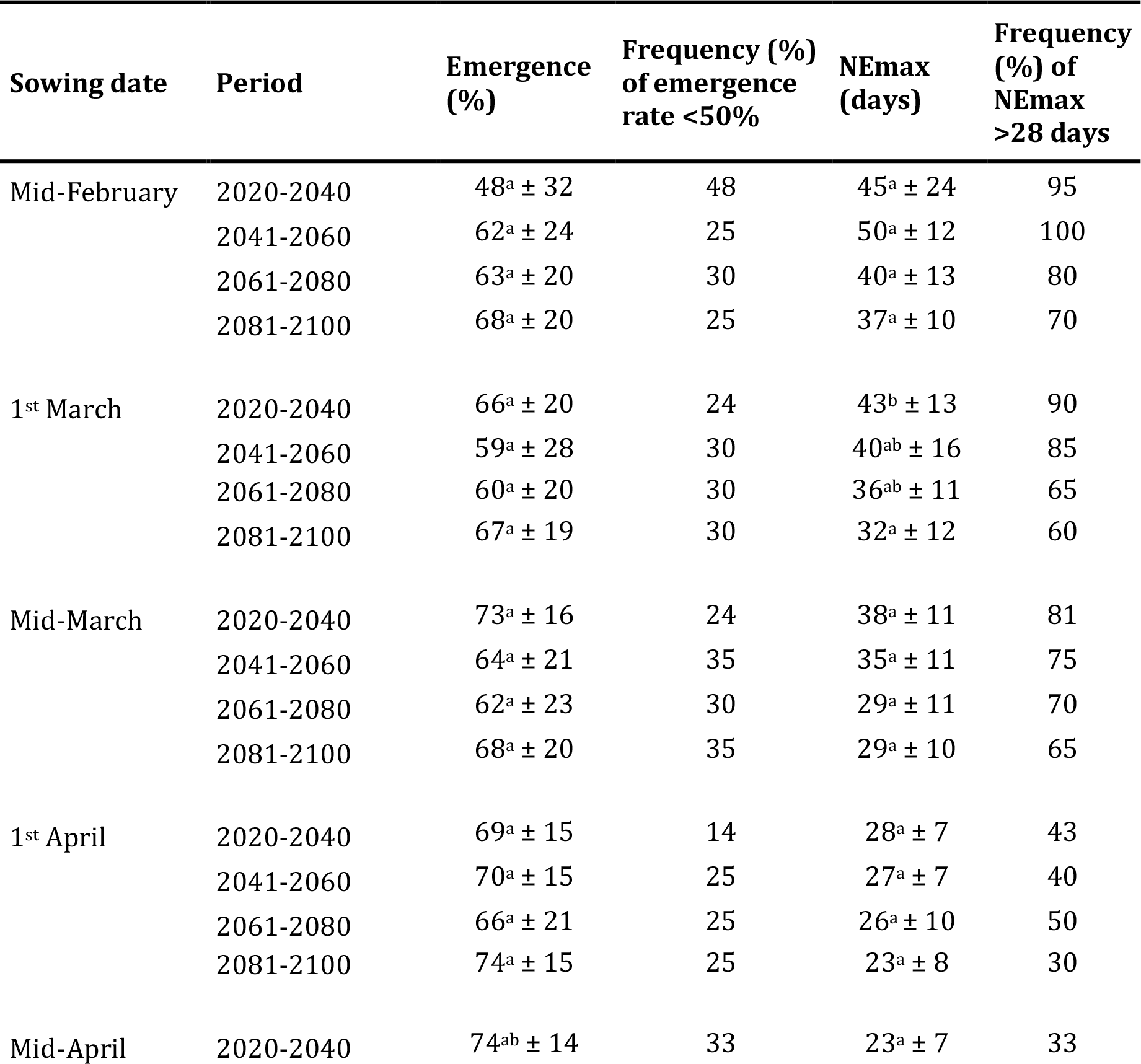
Emergence rate and duration (means ± standard deviation) and frequencies with <50% emergence rate and >28 days to reach the maximum emergence when analyzed by sowing date for each 20-year period and their interaction

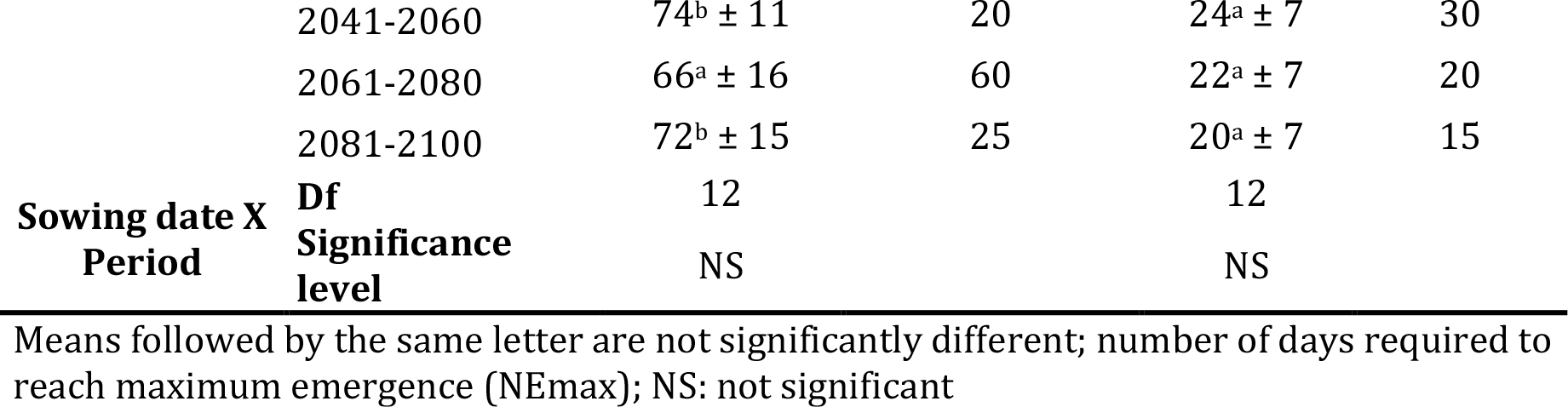

Year-to-year variability in emergence rate was very high for all sowing dates over the 81 simulated years ranging from 0 to 85%. Emergence rate most often registered between 50 and 85%, but emergence rate <50% were observed in many cases. Simulated average emergence rate by “sowing date × period” ranged from less than 50% for the mid-February sowing in 2020-2040 to more than 70% for different sowing dates. The frequency of poor emergence rate (<50%) ranged from 14 to 48% depending on sowing date × period. The mid-February sowing in the 2020-2040 period not only had the lowest average emergence rate but also a high frequency of poor emergence rate.

When data were analyzed by sowing date for each period, the interaction effect of sowing date × period on emergence rate was not statistically significant (P = 0.08). In contrast, the interaction effect was statistically significant (P < 0.001) when all data were pooled and analyzed only by sowing date or period **(Supplementary Table 1)**.

Simulated mean NEmax by sowing date × period ranged from 20 days for the mid-April sowing in the 2081-2100 period to 50 days for the mid-February sowing in the 2041-2060 period **(Table 3)**. Mean NEmax decreased over sowing dates and also over periods, by more than one week for the earlier sowing dates and to a lower extent for later sowings. Within each period, mean NEmax was almost no significantly different between the five sowing dates. However, there were statistically significant differences when the data were analyzed only by sowing date (P < 0.001) or period (P < 0.001**; Supplementary Table 1)**. The frequency of high NEmax (>28 days) ranged from 15 to 100% by sowing date × period. This frequency was higher for earlier sowing dates and also for earlier periods **(Table 3)**.

### 3.4. Causes of non-emergence rates and frequencies

Results on the main causes of non-emergence are reported in **Table 4**. The major causes of non-emergence were non-germination, followed by soil surface crusting and seedling death due to drought while seedling death due to clod was the least important. Results on the effect of sowing date for each period and their interactions on germination rate, duration and frequency are reported in **Table 5**. Outcomes on the overall effect of sowing date and period, and their interactions on germination rate and duration are presented in **Supplementary Table 1**.

**Table 4.**
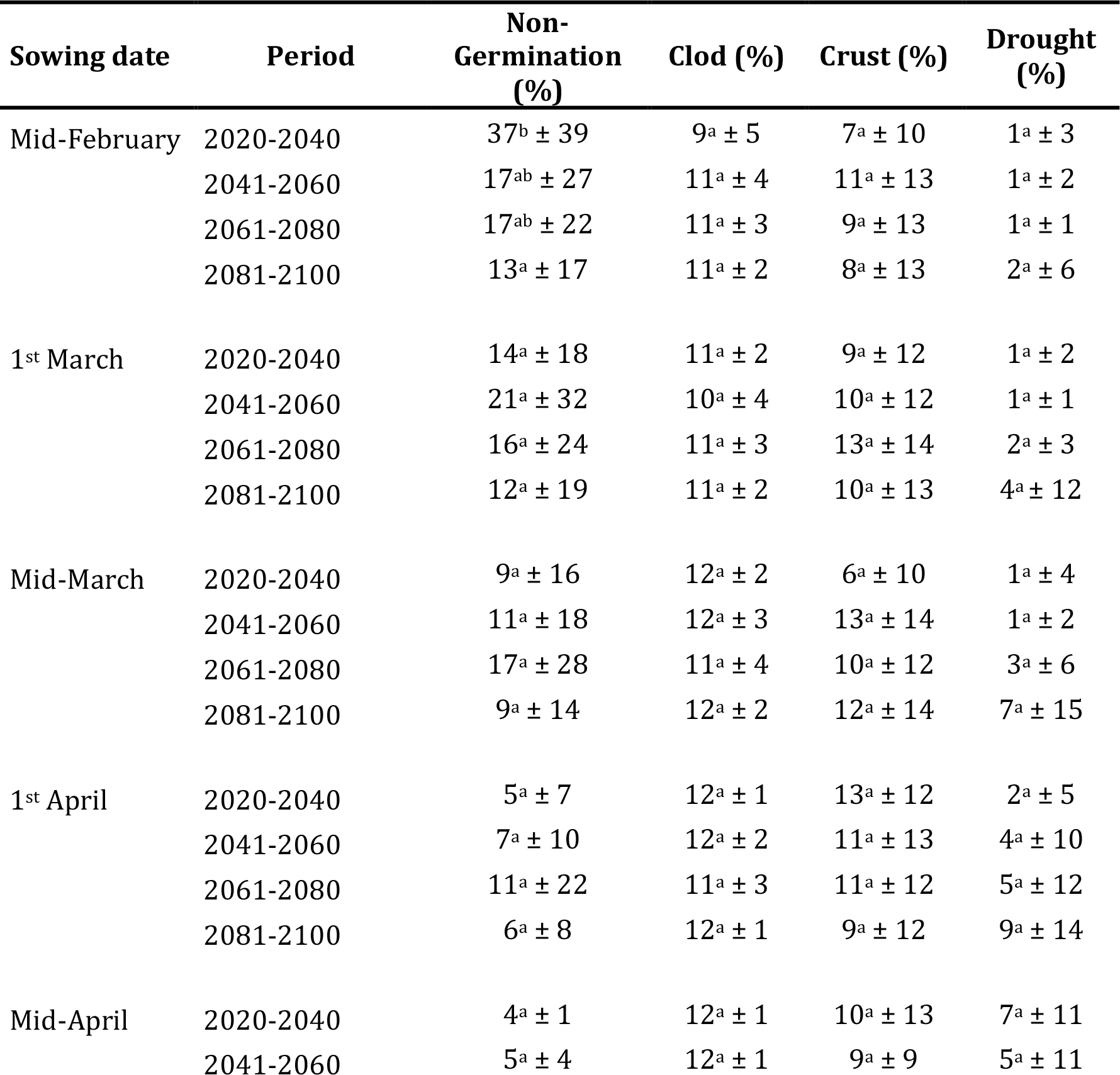
Rates of non-emergence causes (means ± standard deviation) of seedlings as analyzed by sowing date for each 20-year period and their interaction

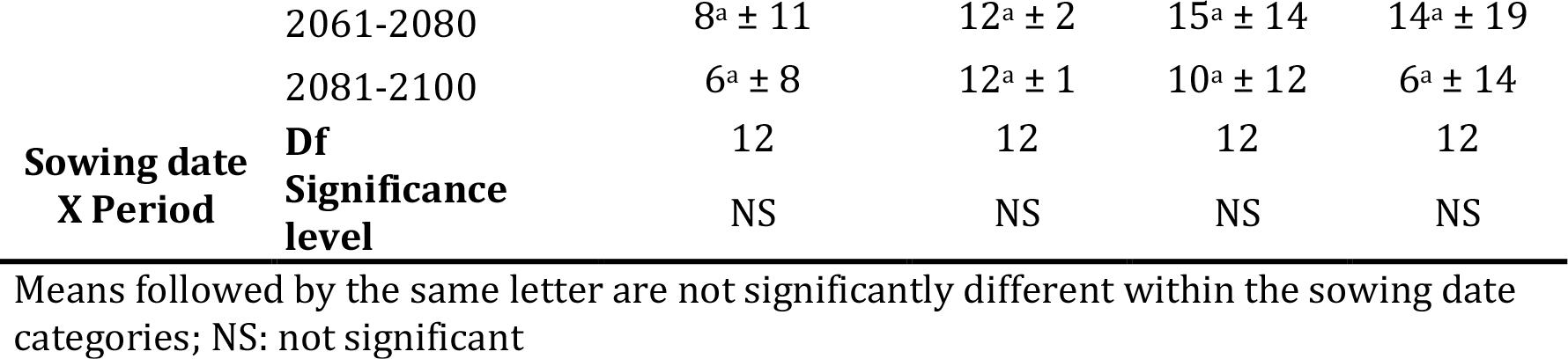

**Table 5.**
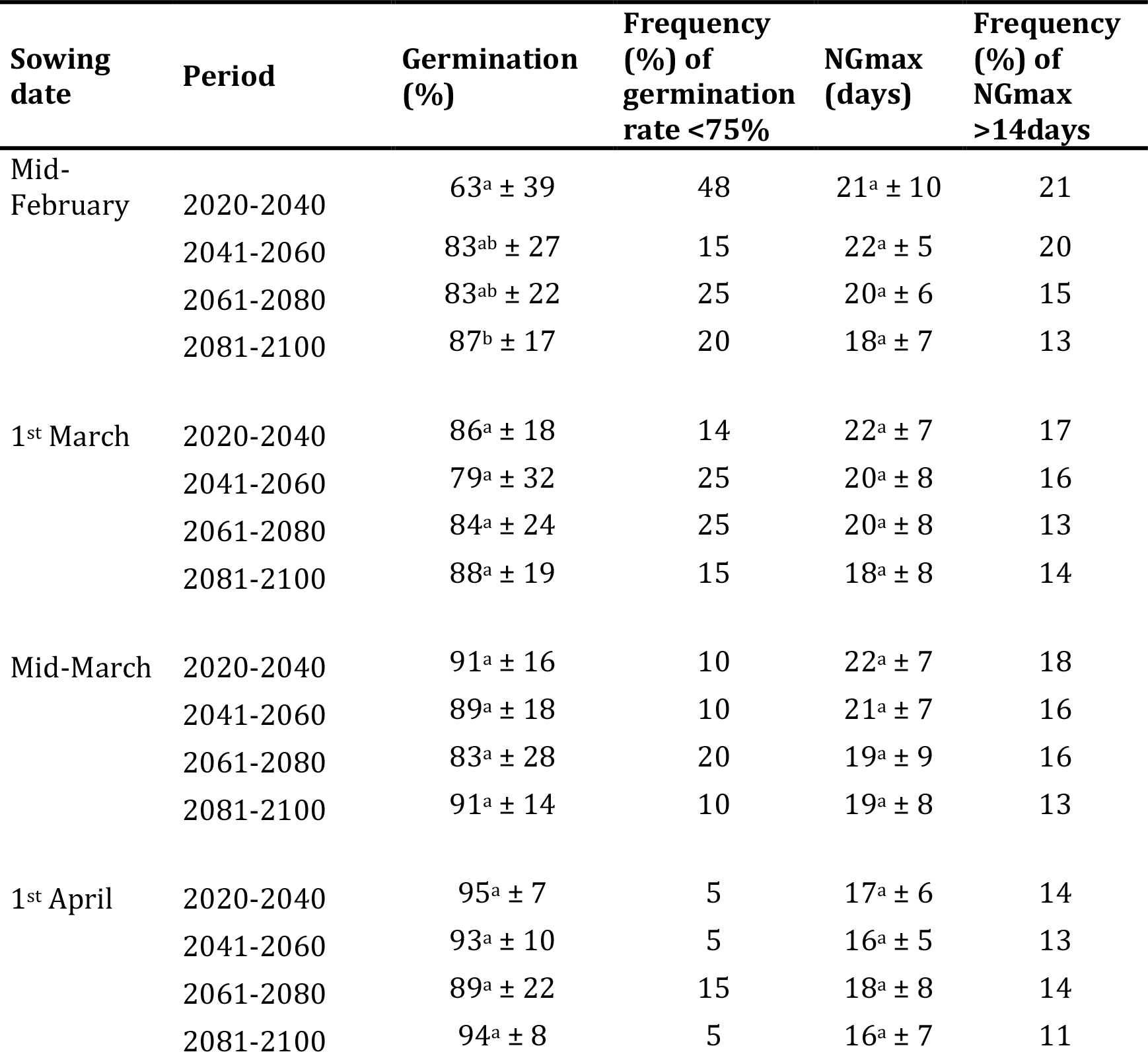
Germination rate and duration (means ± standard deviation) and frequencies with <75% germination rate and >14 days to reach the maximum germination when analyzed by sowing date for each 20-year period and their interaction

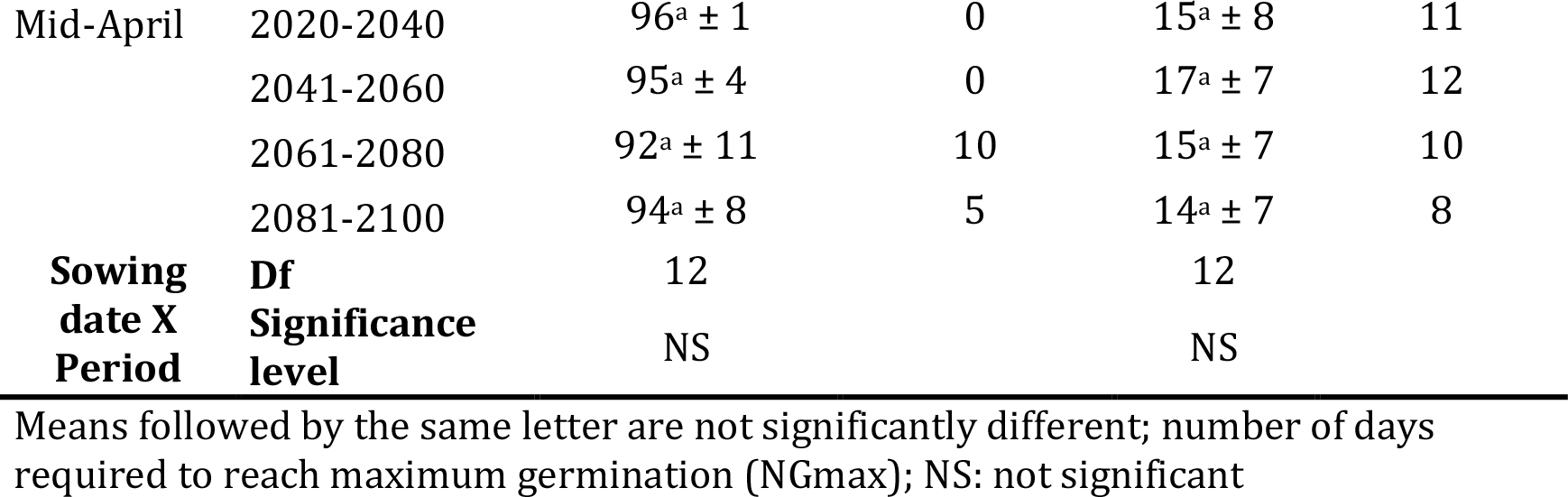

#### 3.4.1. Non-germination

The mean non-germination rate ranged from less than 5% for several sowing dates after the mid-March sowing to 37% for the mid-February sowing in the 2020-2040 period. When data were analyzed by sowing date for each period, and their interaction, no statistically significant effect of the sowing date, period, or their interaction was found on non-germination rate except between the 2020-2040 and 2081-2100 periods for the mid-February sowing. In contrast, non-germination rate differences were statistically significant when the data were combined and analyzed by sowing date (P < 0.001), period (P < 0.01), and their interaction (P < 0.001; **Supplementary Table 2)**. The frequency of high non-germination (>25%) ranged from 0 to 48% **(Figure 3)** depending on sowing date × period and was higher for earlier sowings.

**Figure 3.**
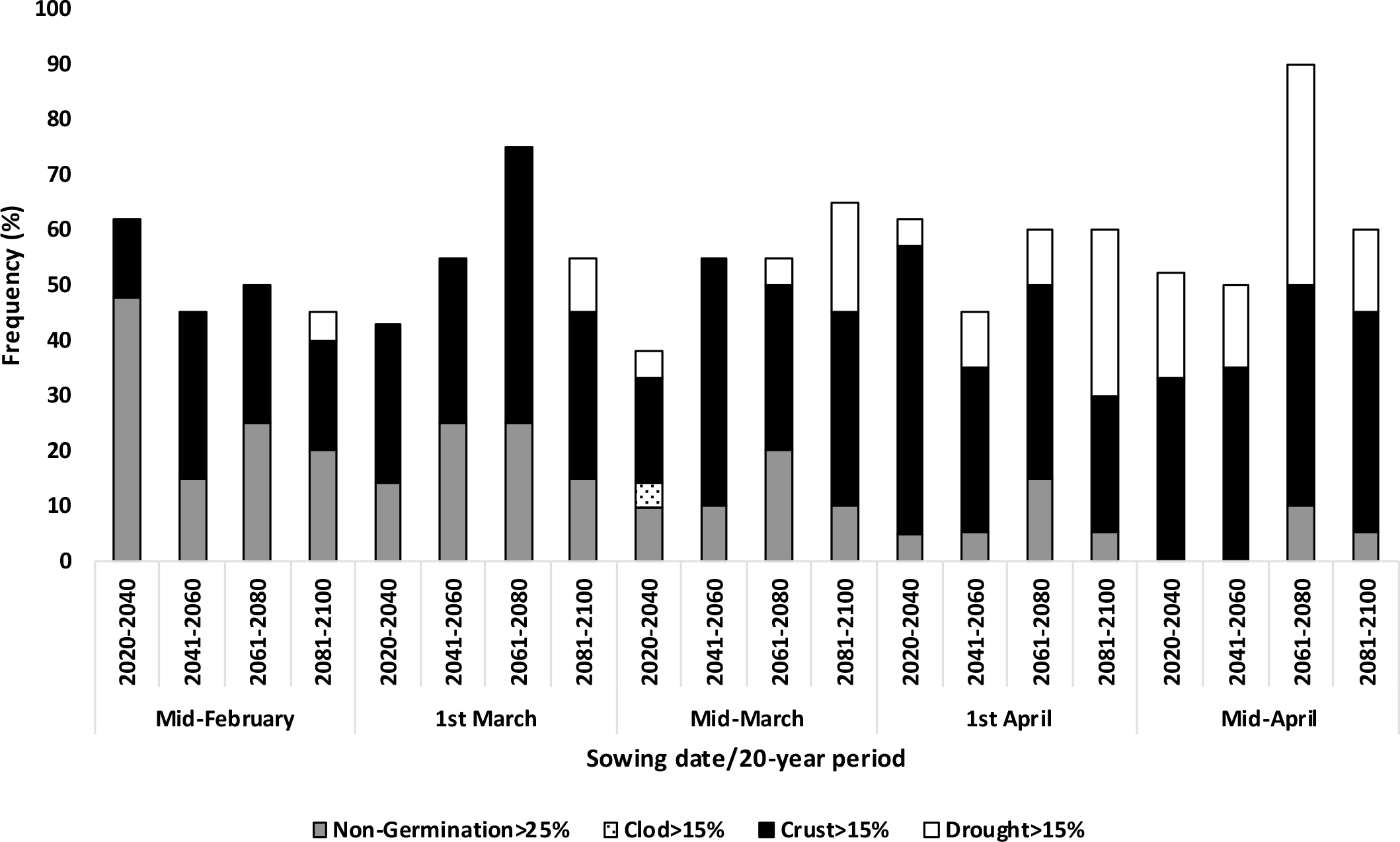
Frequencies (%) of non-emergence causes when analyzed by sowing date for each 20-year period. Only causes with a high frequency that could pose risks of crop emergence failure were considered which included frequency of non-germination >25% and frequency of seedling mortality due to clod, crust and drought, each >15%

Simulated average NGmax values ranged from 14 to 22 days when data were analyzed by sowing date for each period. These values generally decreased with later sowing dates and periods **(Table 5)**. The frequency of high NGmax (>14 days) ranged from 8 to 21% which generally decreased with later sowings and over the periods. No statistically significant effect of sowing date, period and their interaction (P = 0.98) was found on mean NGmax values when data were analyzed by sowing date for each period. In contrast, when data were analyzed only by sowing date or period, there were statistically significant effects of sowing date (P < 0.001) and period (P < 0.01), but not of their interaction on mean NGmax **(Supplementary Table 1)**.

#### 3.4.2. Seedling mortality due to crust

Seedling mortality rate due to soil surface crust ranged from 6 to 15% **(Table 4)**. Average mortality rate was generally lower for the 2020-2040 period until the mid-March sowing, as soil surface crust prevents emergence only when it becomes dry. No statistically significant effect of sowing date (P = 0.48) or period (P = 0.24) or their interaction was observed on seedling mortality rate either when the data were analyzed by sowing date for each period (P = 0.76) or only by sowing date and period (P = 0.22) **(Supplementary Table 2)**. The frequency of high seedling mortality rate (>15%) ranged from 19 to 45% **(Figure 3)**. This frequency was lower for the 2020-2040 period until mid-March sowing.

#### 3.4.3. Seedling mortality due to drought

Seedling mortality rate due to drought ranged from 1 to 14%, and increased with later sowing dates and periods, with some exceptions **(Table 4)**. When data were analyzed by sowing date for each period, no significant effect of sowing date, period or their interaction (P = 0.276) was found on seedling mortality rate. In contrast, when the data were pooled and analyzed only by sowing date and period, statistically significant effect of sowing date (P < 0.001), period (P < 0.001) and their interaction (P < 0.05) was found on seedling mortality rate **(Supplementary Table 2)**. The frequency of high mortality due to drought (>15%) ranged from 0 to 40% **(Figure 3)**. This frequency increased with later sowing dates and periods. It is however remarkable that seedling mortality due to drought appeared for the 2081 – 2100 period even for sowings as early as mid-March or even before.

#### 3.4.4. Seedling mortality due to clod

Seedling mortality rate due to clod ranged from 9 to 12% **(Table 4)** with little variability among the sowing dates or periods. This was expected because this mortality mostly depends on seedbed structure, which was the same for all simulations, independent of the sowing date and period.

### 3.5. Risks of bolting

When data were analyzed by sowing date for each 20-year period, significant effect of sowing date, period or their interaction (P < 0.001) was found on potential bolting rate and devernalization conditions. The average predicted potential bolting rates ranged from 0.04% to 1.65%. As expected, bolting rates were higher for sowings in February and decreased with later sowing dates. Our results showed that the predicted bolting rates decreased progressively and significantly after 2060 for all simulated sowing dates **(Table 6**). Likewise, the potential for devernalization highly increased due to an increased number of days with Tmax > 25°C at the end of spring **(Table 6)**. Based on these results, the average risk of bolting will be lower after 2060, even for the earliest sowing dates.

**Table 6.**
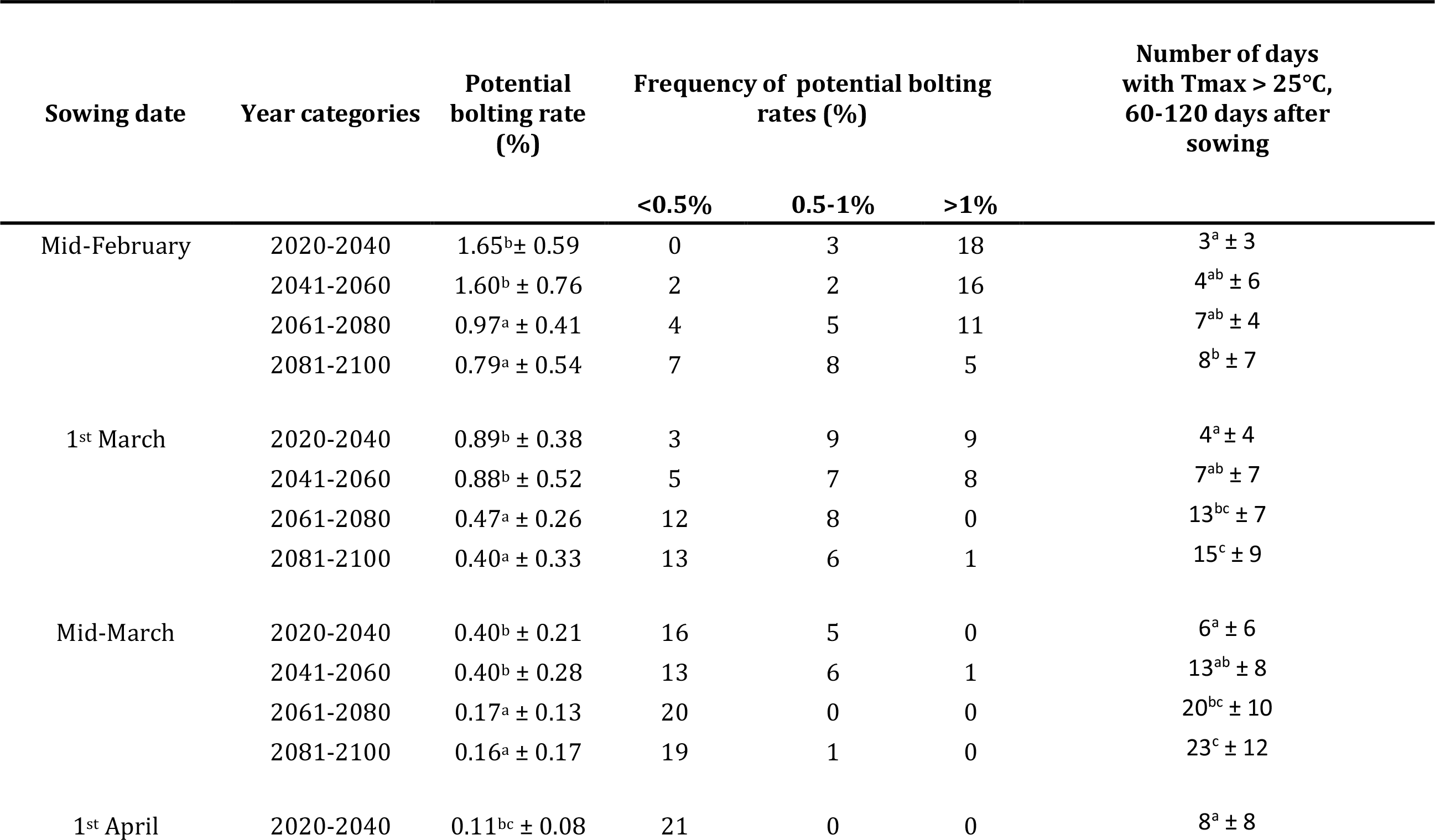
Bolting rate (means ± standard deviation), without the devernalization effect, and frequency when analyzed by sowing date for each 20-year period and their interaction and potential devernalization due to high temperatures (7 days with Tmax > 25°C) 60 to 120 days after sowing.

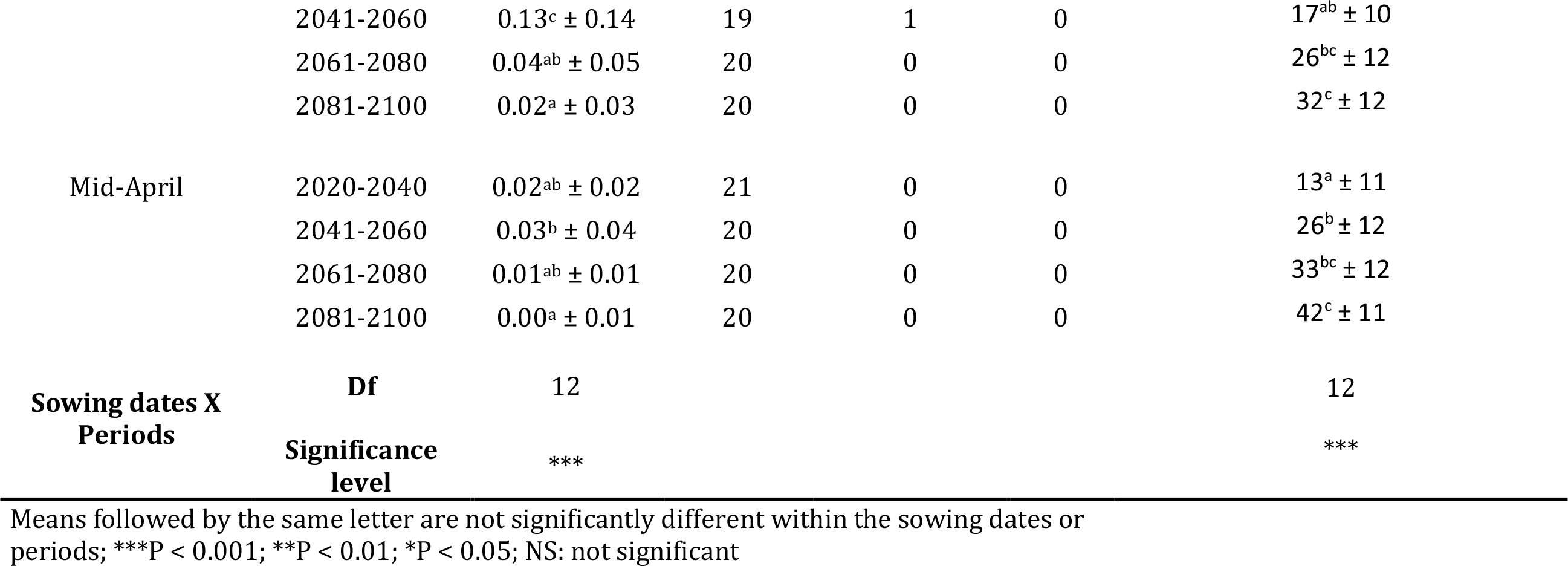

### 3.6. Technical feasibility of sowing

Results on the probability of field access for February and March over periods are reported in **Table 7**. When considering cumulated rainfall over one month or the number of days >1 mm rainfall in February or March (i.e. the earliest sowing periods), there were no statistically significant differences between the two months and over the 20-year periods. This means that the technical feasibility of sowings will remain the same as nowadays, as shown in **Figure 1**.

**Table 7.** Indicators for field accessibility for farmers to perform sowing for early sowing dates and periods

| Period of time | Period | Average cumulated rainfall<br>30 days before sowing (mm) | Number of days with<br>>1mm |
| --- | --- | --- | --- |
| 1<sup>st</sup> February – end of February | 2000-2018 | 38<sup>a</sup> ± 3 | 8<sup>a</sup> ± 0.45 |
|  | 2020-2040 | 47<sup>a</sup> ± 3 | 11<sup>a</sup> ± 0.47 |
|  | 2041-2060 | 51<sup>a</sup> ± 3 | 11<sup>a</sup> ± 0.48 |
|  | 2061-2080 | 51<sup>a</sup> ± 4 | 10<sup>a</sup> ± 0.47 |
|  | 2081-2100 | 52<sup>a</sup> ± 3 | 11<sup>a</sup> ± 0.47 |
| 1<sup>st</sup> March –end of March | 2000-2018 | 45<sup>a</sup> ± 3 | 8<sup>a</sup> ± 0.43 |
|  | 2020-2040 | 48<sup>a</sup> ± 3 | 10<sup>a</sup> ± 0.45 |
|  | 2041-2060 | 50<sup>a</sup> ± 3 | 9<sup>a</sup> ± 0.44 |
|  | 2061-2080 | 37<sup>a</sup> ± 3 | 7<sup>a</sup> ± 0.42 |
|  | 2081-2100 | 50<sup>a</sup> ± 4 | 9<sup>a</sup> ± 0.45 |

Means followed by the same letter are not significantly different within the sowing dates or periods

## 4. Discussion

### 4.1. Seedbed micro-climatic conditions under future scenarios

We used the STICS soil-crop model based on the RCP 8.5 emission scenario to generate climate data. This model has been reported to be sensitive enough to generate realistic soil data such as soil moisture (Constantin et al. 2015; Dürr et al. 2016; Tribouillois et al. 2018). Predictions of the STICS soil-crop model showed that during the sowing period of sugar beet, mean seedbed temperatures will increase over time and that a higher variability of rainfall will occur, without an overall increase of its cumulated values. The trends we found here are coherent with those generally expressed for global climate changes, and the use of the STICS model allows to evaluate more precisely these changes specifically in the seedbed and for the sowing period of sugar beet in Northern France or Europe.

Under the most pessimistic climate scenario that we used, the predicted rise in mean soil temperature at sowing remained 0 °C until 2060 and became +2 °C after 2080. When climatic data of the next eight decades were compared with the past two decades, we found that average soil temperature 30 dps of the last 19 years were similar to those predicted until 2060, but higher over the last two periods. This highlights that the impact of climate change will become more remarkable after 2060 with warmer soil temperatures during the last two decades of the 21^st^ century. Interestingly, when maximum air temperature at sowing was considered, mean values 30 dps showed a very high year-to-year variability, but without any regular increase over the years.

In contrast to predicted seedbed temperatures, cumulative rainfall did not change over time and were more or less the same for the first four sowing dates. A delay of two weeks in sowing from 1^st^ to mid-April resulted in an increased drought risk under future climate change.

### 4.2. Sugar beet crop establishment under future climate

Several previous studies compared results of field observation and simulation using the SIMPLE crop emergence model and found its prediction similar to observed data (Dorsainvil et al. 2005; Brunel-Muguet et al. 2011; Constantin et al. 2015; Dürr et al. 2016). Therefore, prediction of germination and emergence rates reported in this study can be considered reliable. Even by using the most pessimistic climate scenario, predictions based on the SIMPLE crop emergence model showed that, in most cases, there will be a sufficient level of sugar beet crop emergence in Northern France and Europe under future climate change.

Despite performing simulation studies using only one climate scenario, the results of this study represent an important outcome for decision making related to sugar beet sowing not only in Northern France, but in Northern Europe in general due to similar climatic conditions and sowing dates. The inclusion of the most pessimistic climate scenario for simulation did not render necessary the use of other less pessimistic climate scenarios (i.e. RCP 2.6, 4.5 and 6). This is because we did not find any dramatic changes in sugar beet emergence rate which would have been less impacted with simulation studies including less drastic future climate scenarios. Nevertheless, our results are based on only one study site which represents a limit and thus future studies taking into account several study sites over space could shed more light in this regard.

The most important finding of this study is that there are no important variability in terms of emergence rate among sowing dates, except for the earliest one for which emergence rate was predicted to be higher and less variable after 2060. Sowing date adaptation is, by far, the most frequently investigated climate change adaptation option (White et al. 2011). Sugar beet farmers in France and Northern Europe, who currently practice the mid-March sowing, may thus anticipate sowing under future climate scenarios, given that earlier sowing provides higher yield benefits. This is due to a prolonged vegetation period and the higher amount of intercepted solar radiation, as it is the case for many field crops (Van Ittersum and Rabbinge 1997).

Bolting causes yield penalties in sugar beet, and contribute to gene flow, seed dispersion, and volunteer plant development in the next crops (Longden et al 1975; Sester et al 2008). Therefore, bolting risks could be a limiting factor even when there are possibilities for earlier sowing. Our results showed that the predicted bolting risk will decrease over time and will become reduced for the mid-February sowing and very limited for the 1^st^ March sowing, especially after 2060.

### 4.3. Causes of non-emergence of seedlings under future scenarios

In terms of the total percentage of emergence failure, the one due to non-germination was the most important followed by soil surface crusting and drought. Seedling mortality rates under clod did not vary over sowing dates or periods since it strictly depends on the seedbed structure chosen for simulations. It is also the reason why the maximum simulated emergence rate remained always around 85%, due to about 5% non-germinating seeds in the simulated seed lot and about 10% non-emerging seedlings due to the simulated seedbed structure. Both germination and emergence were affected by the considered abiotic stresses. At the germination stage, very low temperature with earlier sowings, and very low or no rainfall during the later sowings affected the seed germination process. During the emergence phase, the frequency of emergence failure was either related to seedling mortality due to a soil surface crust with all sowing dates, or to water stress with later sowings. The average risk of crop emergence failure remains similar with sowing dates or periods but the prevalence of individual stress factor changes according to sowing dates and periods. After 2060 and to a greater extent after 2080, higher risks of seedling mortality due to drought appear even for the earliest sowing date. Such an analysis of non-emergence results can be obtained only with a simulation approach. Even in the current situation, field observations are rarely undertaken since they are difficult, time consuming and cannot be performed in a high number of fields.

Although the SIMPLE model does not consider the effect of high temperatures that could inhibit germination, we exclude the impact of this stress, given that all sowing were performed in spring and under North European conditions.

Seed germination and seedling emergence rates of sugar beet simulated by the SIMPLE crop emergence model could be overestimated because this model does not take into account the effect of biotic stresses (Constantin et al. 2015). Nevertheless, the risk related to biotic stress could be still limited under current cropping practices for two reasons. First, pelleting of sugar beet seeds containing protectants (fungicides, insecticides, and nematicides) and biostimulants -- is performed to date on 100% seeds (Agreste 2014) which may limit risks of the sugar beet crop establishment due to biotic stresses. Although several diseases of sugar beet caused by soil-borne pathogens, including Rhizoctonia root rot and damping-off, have been reported in Northern France (Motisi et al. 2009), the disease pressure is generally low when seeds are treated. Secondly, sugar beet crop is often rotated with other crops including wheat, to reduce pest inocula *sensu lato*, although some of the crops introduced into the rotation scheme may also be affected by the same soil-borne pathogens affecting sugar beet (Motisi et al. 2009). This is due to a wide host range of most soil-borne pathogens affecting the crop establishment phase (Lamichhane et al. 2017). Therefore, risks related to biotic stresses may be a limiting factor to sugar beet crop establishment under two conditions: i) when seeds are not treated with conventional pesticides and when farmers plan to anticipate sowing, especially under climate change. As shown in this study, an anticipation of sowing, compared to the currently practiced sowing (i.e. mid-March) may be beneficial in terms of yield, but it has to take into account potential risks due to biotic stresses. The latter is generally increased when crops are sown into cold and humid soil conditions and without chemical seed treatment (Serrano and Robertson 2018). Therefore, future studies should integrate the biotic determinants affecting crop establishment into the SIMPLE crop emergence model since the sustainability of chemical pesticides in general and those used for seed treatment in particular is increasingly questioned, especially in the European Union for human health and environmental reasons (Lamichhane et al. 2016). This has led to the recent ban of neonicotinoids in the EU which were widely used for seed treatment (Gross 2013).

### 4.4. Technical feasibility of sowing

The feasibility of technical field operations depends on water content of the soil top layers and thus also on climate change and sowing dates. We evaluated the possibility to enter into the field with agricultural equipments including a seeder for each simulated sowing date and year, using past historical data on earlier sowing dates. Our results suggest that, field access will represent the main limit for earlier sowings in the future as rainfall during early spring will not decrease, compared with the past.

## 5. Conclusions

Climate impact studies are dominated by those on crop yields (Wollenberg et al. 2016). Little is known about the impact of changing climate on specific stages of the crop cycle, especially the crop establishment phase. To achieve an acceptable level of yield it is essential to optimize conditions that favor crop establishment. Despite several limitations, simulation studies represent an important means when it comes to predict food security of the 21^st^ century under future climate change. The present study provides important information that was not possible without mobilizing simulation approach using process-based models. Despite some possibilities of crop emergence failure, the quality of crop establishment will be acceptable under future scenarios, which was not easy to predict without simulations. An anticipation of sowing, compared to the currently practiced sowing (i.e. mid-March), will be viable under future climate change, with possibility of compensating increasing drought risks during summer. However, the possibility of filed access will remain a limiting factor due to extremely variable and high cumulative rainfall values in late winter across our study sites.

## Acknowledgements

This study was supported by a starter grant of the INRA’s Environment and Agronomy Division to the first author. The authors thank the colleagues from technical institute of sugar beet (ITB) for useful discussion on this topic.

---

**Supplementary Table 1.** Germination and emergence rates and duration of sugar beet (means ± standard deviation) when analyzed by sowing date and 20-year period

| **Sowing date** | **Germination (%)** | **Emergence (%)** | **NGmax (days)** | **NEmax (days)** |
| --- | --- | --- | --- | --- |
| Mid-February | 79<sup>a</sup> ± 29 | 59<sup>a</sup> ± 25 | 17<sup>a</sup>±7 | 26<sup>b</sup>±8 |
| 1<sup>st</sup> March | 84<sup>ab</sup> ± 23 | 61<sup>a</sup> ± 22 | 20<sup>b</sup>±7 | 38<sup>d</sup>±14 |
| Mid-March | 89<sup>bc</sup> ± 19 | 64<sup>a</sup> ± 21 | 15<sup>a</sup>±7 | 22<sup>a</sup>±7 |
| 1<sup>st</sup> April | 93<sup>c</sup> ± 13 | 64<sup>a</sup> ± 20 | 20<sup>b</sup>±7 | 43<sup>e</sup>±16 |
| Mid-April | 94<sup>c</sup> ± 7 | 63<sup>a</sup> ± 20 | 20<sup>b</sup>±8 | 33<sup>c</sup>±11 |
| Df | 4 | 4 | 4 | 4 |
| Significance level | *** | NS | *** | *** |
| **Period** |  |  |  |  |
| 2020-2040 | 86<sup>a</sup> ± 23 | 63<sup>ab</sup> ± 23 | 19<sup>b</sup>±8 | 35<sup>b</sup>±16 |
| 2041-2060 | 88<sup>a</sup> ± 21 | 63<sup>ab</sup> ± 21 | 19<sup>b</sup>±7 | 35<sup>b</sup>±14 |
| 2061-2080 | 86<sup>a</sup> ± 22 | 58<sup>a</sup> ± 22 | 18<sup>ab</sup>±8 | 31<sup>a</sup>±12 |
| 2081-2100 | 91<sup>a</sup> ± 14 | 64<sup>b</sup> ± 21 | 17<sup>a</sup>±7 | 29<sup>a</sup>±11 |
| Df | 3 | 3 | 3 | 3 |
| Significance level | NS | * | ** | *** |
| **Sowing date X Period** |  |  |  |  |
| Df | 12 | 12 | 12 | 12 |
| Significance level | *** | *** | NS | NS |

Means followed by the same letter are not significantly different within the year or sowing date categories; \*\*\*P < 0.001; \*\*P < 0.01; \*P < 0.05; NS: not significant; NGmax and NEmax: number of days required to reach the maximum germination and emergence respectively

**Supplementary Table 2.** Causes of non-emergence rates of sugar beet (means ± standard deviation) when analyzed by sowing date and 20-year period

| **Sowing date** | **Non-Germination (%)** | **Clod (%)** | **Crust (%)** | **Drought (%)** |
| --- | --- | --- | --- | --- |
| Mid-February | 7<sup>a</sup> ± 13 | 10<sup>a</sup> ± 4 | 9<sup>a</sup> ± 12 | 2<sup>a</sup> ± 4 |
| 1<sup>st</sup> March | 16<sup>bc</sup> ± 23 | 11<sup>ab</sup> ± 3 | 10<sup>a</sup> ± 12 | 3<sup>a</sup> ± 6 |
| Mid-March | 6<sup>a</sup> ± 7 | 11<sup>bc</sup> ± 3 | 10<sup>a</sup> ± 12 | 4<sup>ab</sup> ± 8 |
| 1<sup>st</sup> April | 21<sup>c</sup> ± 29 | 12<sup>c</sup> ± 2 | 11<sup>a</sup> ± 12 | 6<sup>bc</sup> ± 11 |
| Mid-April | 11<sup>ab</sup> ± 19 | 12<sup>c</sup> ± 1 | 11<sup>a</sup> ± 12 | 9<sup>c</sup> ± 14 |
| Df | 4 | 4 | 4 | 4 |
| Significance level | *** | *** | NS | *** |
| **Period** |  |  |  |  |
| 2020-2040 | 14<sup>a</sup> ± 23 | 11<sup>a</sup> ± 3 | 9<sup>a</sup> ± 12 | 3<sup>a</sup> ± 7 |
| 2041-2060 | 12<sup>a</sup> ± 21 | 11<sup>a</sup> ± 3 | 10<sup>a</sup> ± 12 | 3<sup>a</sup> ± 7 |
| 2061-2080 | 14<sup>a</sup> ± 22 | 11<sup>a</sup> ± 3 | 11<sup>a</sup> ± 13 | 6<sup>b</sup> ± 12 |
| 2081-2100 | 9<sup>a</sup> ± 14 | 11<sup>a</sup> ± 2 | 10<sup>a</sup> ± 12 | 6<sup>b</sup> ± 12 |
| Df | 3 | 3 | 3 | 3 |
| Significance level | ** | NS | NS | *** |
| **Sowing date X Period** |  |  |  |  |
| Df | 12 | 12 | 12 | 12 |
| Significance level | *** | *** | NS | * |

Means followed by the same letter are not significantly different within the year or sowing date categories; \*\*\*P < 0.001; \*\*P < 0.01; \*P < 0.05; NS: not significant

